# Effects of a transmissible cancer on life-history traits in Tasmanian devils

**DOI:** 10.1101/2024.10.14.618139

**Authors:** Kasha Strickland, Menna Jones, Shelly Lachish, Sebastien Comte, Rodrigo Hamede, Paul Hohenlohe, Hamish McCallum, Andrew Storfer, Loeske Kruuk

**Affiliations:** Institute of Ecology and Evolution, School of Biological Sciences, University of Edinburgh, UK; School of Natural Sciences, University of Tasmania, Hobart, TAS, Australia; Public Health Intelligence Branch, Queensland Public Health and Scientific Services Division, Queensland Health, 15 Butterfield Street, Herston, QLD 4006; Vertebrate Pest Research Unit, NSW Department of Primary Industries, 1447 Forest Rd, Orange NSW 2800, Australia; Department of Biological Sciences, University of Idaho, Moscow, Idaho 83844; Environmental Futures Research Institute, Griffith University, Nathan, Queensland, Australia; School of Biological Sciences, Washington State University, Pullman, Washington, USA 99164-4236

**Keywords:** life history traits, phenotypic plasticity, precocial breeding, inbreeding depression

## Abstract

Shifts in life history traits, such as timing of reproduction, can help mediate population declines following perturbations, and early reproduction should be favoured when adult survival is impacted more than juvenile survival. In Tasmanian devils, following the emergence of a fatal transmissible cancer, females started to breed precocially (i.e., at age one instead of typically age two) and the same time as populations started to decline following disease emergence. Here, we focus on a diseased site (Freycinet Peninsula, Tasmania, Australia) with 18 years of continuous mark-recapture data to test: (1) whether rates of precocial breeding in females continued to increase after the initial rise after the emergence of the disease, (2) whether there was a relationship between body size and breeding success for either one-year-olds or adult females (i.e., at least two-years-old), and (3) whether there was inbreeding depression in breeding success for either age category. We show that rates of precocial breeding did not continue to rise, and that the proportion of precocially breeding females has plateaued at around 50%. We also show that there was no effect of body size on the probability of breeding for either one-year-old or for adult females. Finally, we show that there was no evidence for inbreeding depression in breeding success for either age class. We discuss possible constraints that may have inhibited further rise in rates of precocial breeding in the context of limitations to growth in the offspring of precocially breeding (and therefore smaller) females.

## Introduction

One of the most fundamental trade-offs in life-history theory involves that between growth and timing of reproduction (1). Specifically, resource limitations experienced in wild populations results in a need for individuals to differentially allocate energy to competing life-history functions (2,3). Given that the resultant trade-off is often associated with shifts in allocation to different vital rates (i.e., survival vs reproduction), life-history strategies adopted by populations can have important effects on population dynamics (4–6). Consequently, shifts in life-history traits can be one of the first responses to population perturbances (either via plasticity or evolution) and, in some cases, can mitigate their impacts on population decline (7– 9). For instance, it has been shown that maturation and the timing of reproduction may become earlier following population declines caused by shifts in predation (10–12), over-fishing (13,14), or disease outbreaks (15,16).

While shifts in life-history traits may buffer the effect of extrinsic drivers of population decline (17), another important impact is the risk of inbreeding depression (18), which can vary in strength across the lifespan (19). Inbreeding depression describes a decline in fitness-related traits caused by the expression of deleterious recessive alleles as a result of increased homozygosity resulting from mating between relatives (20). As populations decline, the potential for inbreeding increases (21), and coupled with the decline in genetic diversity that is usually associated with population decline (22), inbreeding depression can be a major risk for declining populations. Further, the strength of inbreeding depression may vary with age (19,23,24). For instance, the mutation accumulation hypothesis suggests that inbreeding depression may be stronger in later life because selection fails to purge deleterious alleles which are expressed in later life (25). Alternatively, increased mortality of inbred individuals at early life stages may result in decreased inbreeding depression in later life stages (24). It is therefore important to consider how inbreeding depression differentially affects life-stages when studying how life-history traits might shift following perturbations to populations.

One important example of a life-history shift during population declines has been in Tasmanian devils, where females started to breed precocially following the emergence of a fatal, transmissible cancer (15). Devil facial tumour disease (DFTD) has caused declines of up to 80% across the geographic range of Tasmanian devils, which are the largest carnivorous marsupial endemic to Tasmania, Australia (26,27). Prior to the emergence of DFTD, almost all female devils reached sexual maturation and usually started to breed at two years of age (15,28). However, in the two years immediately after the emergence of the disease, approximately 40% of females began to breed at one year of age (15). Such “precocial” breeding is thought to have arisen due to increased food availability associated with the decreasing population sizes, which would have relaxed the trade-off associated with growing to the threshold size to enable sexual maturity. However, it has also been suggested that the offspring of precocial breeders may not be able to breed precocially. This is because females that breed precocially normally give birth at about 14 months, which is later during the mating season (May versus February-March) than individuals two or more years old. Therefore their offspring are unlikely to reach sexual maturity by May of the following year due to being younger and therefore smaller (28). Consequently, constraints on continued increases in rates of precocial breeding may arise. Further, it follows that larger juvenile females may be more likely to breed earlier, indicating a life-history shift that relaxes the growth-reproduction trade-off. (29).

In this study, we aimed to follow-up on previous studies that showed evidence of a rise in precocial breeding in females (15). We analysed an additional 18 years of data collected as part of a long-term mark-recapture study of Tasmanian devils on the Freycinet peninsula, Tasmania. Specifically, we tested: (1) whether the initial increase in rates of precocial breeding following emergence of DFTD continued over time, (2) whether there was evidence for a relationship between age at breeding and size (measured as both body weight (kg) and head width (mm)), and (3) for evidence of inbreeding depression in annual breeding success for both 1-year olds and for adults (i.e., at least 2-years-old).

## Materials and Methods

### Tasmanian devil study site, trapping and phenotypic data

We used mark-recapture data collected on Tasmanian Devils from Freycinet Peninsula, Tasmania, Australia between January, 1999 and May, 2021. Tasmanian devils were trapped across the entire peninsula up to four times a year using custom-built baited traps (30), with trapping periods timed to coincide with key stages in the breeding cycle: autumn (March/April), early pouch young; winter (June/ July), late pouch young; spring (September/October), females lactating with young in dens; summer (December/January), dependent young emerging from dens before weaning in early February. During their first capture, devils were sexed, individually tagged with a microchip and had a tissue sample taken for genetic analysis (see below). At first capture and then at all subsequent recaptures, their age, head width (in mm; precise linear measure of body size taken across the jugal arch) and body mass (in kg) were recorded as described in (30). Individuals were aged using a combination of head width, molar eruption, molar tooth wear and canine over-eruption (31), and given a birthdate of April 1^st^ for a given year as per Lachish et al. (2007). This method of aging is accurate up to two years of age, but most individuals were first trapped as juveniles and are therefore of known age. Disease status (presence/absence) was determined for each capture by visual inspection for tumours and/or histopathological examination of tumour biopsies (32).

We used a subset of these data that included repeated observations of females of known age, and where size and reproductive status had been recorded. For each observation of females, reproductive status was determined using the following criteria: presence of active (lactating) teats, presence of pouch young, and pouch appearance (33). Using these criteria, we measured whether a female had been observed as having bred or not within that year (defined as April - April), which we hereafter refer to as annual breeding success, which was recorded as a binary variable. More than one observation per female in a given year were reduced to a single estimate of breeding success. In cases where status varied among observations we retained just the first observation that confirmed breeding.

We subset the data to include observations of individuals using a minimum age of 14 months. This was done (1) in order to reduce the conflation between age and size measurements, (2) because 14 months is the youngest age at which females have bred in the dataset. Re-analysing the data using an age of 18 months as a threshold did not change the qualitative inferences of the results, but did reduce the dataset such that confidence in parameter estimates decreased. Therefore, we retained the 14-month minimum age cut-off. Age was binned into two age categories for all subsequent analyses: 1-year-old (between 14 – 23 months old), and adults (24 months old or older). Because there were very few observations of 1-year old breeding females prior to 2003, we removed those years from further analyses. This resulted in a dataset which contained a total of 372 observations of 253 females caught between 2003 and 2021 for further analyses.

### DNA extraction and genotyping

We extracted DNA from tissue samples using DNeasy kits (Qiagen), and genotyping was as previously described in (34,35). Briefly, single-nucleotide polymorphism (SNP) genotyping was achieved via single-digest *RADcapture* (i.e., “Rapture” (36) of DNA extracted from tissue. All raw reads from sequencing were first aligned to to the *S. harrisii* reference genome (37). PCR duplicates were removed, and SNP calling was conducted using *gtstacks* (Catchen et al. 2013) on the merged “bam” files from reads generated from two rounds of sequencing (which was done to achieve sufficient read depth (Margres et al. 2018)). The function *populations* was then used to keep one random SNP per RAD locus and per 10Kb window, exclude SNPs with a minor allele frequency (MAF) below 1%, to remove individuals with more than 70% missing data, and to remove SNPs present in less than 50% of the samples. We then further filtered genotype calls with a read depth of <4 to increase genotyping accuracy, before reapplying the filtering parameters explained above. This resulted in a total of 2105 SNPs genotyped in a total of 584 individuals for the whole study population (which was further restricted for use with phenotypic data in our analyses – see below).

### Inbreeding coefficients

We measured variation in inbreeding using genomic inbreeding coefficients estimated in GATK (38). We used 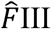 (hereafter F_GRM_), which estimates the allelic correlation between gametes, which is closely correlated with runs of homozygosity on the genome (F_ROH_), and is therefore a reliable measure of the genomic consequences of inbreeding (38). We ensured that F_GRM_ measures were robust to SNP filtering by varying the MAF cut-off criterion at 1%, 5% and 10%. F_GRM_ estimates were all very highly correlated regardless of which MAF cut-off we used (r > 0.99%). F_GRM_ ranged from -0.37 (indicating outbreeding) to 0.36 (indicating significant inbreeding) (median F_GRM_ = -0.04, variance = 0.006).

### Statistical analyses

In all models described below, annual breeding success was fit as a binary variable with a logit link via the Bernoulli family in stan using the *brms* R package (39). Models were fit with normal priors with 5 standard deviations on the fixed effects and half-Cauchy priors with 3 degrees of freedom on the random effects. All models were run for 6000 iterations with a warmup period of 2000 runs across four chains, and convergence was assessed by ensuring R- hat was below 1.01, effective sample sizes for all parameters were at least 1000 and by visually ensuring chains had mixed well.

The first model we ran was used (1) to determine whether there was a temporal trend in probability of precocial breeding, and (2) to estimate the relationship between size (measured as body weight (kg) and head width (mm)) and annual breeding success, and whether this relationship differed between 1-year olds and adults. This model was therefore fit with the following variables included as fixed effects: year, age (defined as a category with two levels: “1-year-olds” or “adults”), head width, body weight and DFTD status (present or absent). DFTD status was included because it affects breeding success (28). We also modelled the interaction between size traits and age to determine whether larger 1-year-old females were more likely to breed than smaller ones. We further fit the interaction between year and age to determine whether there was age-dependent temporal change in the probability of breeding. Year, individual ID, trap ID (i.e., the trap number in which the individual was caught, the locations of which are the same throughout the study period) and calendar month were all included as random effects to account for non-linear temporal changes, repeated measures for individuals, spatial heterogeneity and seasonality of breeding. To test whether there was non- linearity in the effect of year on annual breeding success for either 1-year-olds or adults, we refit the model for annual breeding success described above but instead fit year as a smoothed term with 5 knots. Fitting it with more knots did not change our overall inferences (see supplemental Table 1).

To investigate whether there was evidence for inbreeding depression in annual breeding success, we ran a model of breeding success with age (as a two-level category “1-year-olds” or “Adults”) and F_GRM_ as fixed effects. We further included the interaction between age and F_GRM_ to determine whether inbreeding depression was more prevalent in one age category than the other. As above, year, ID, trap ID and calendar month were included as random effects. This model was run with a smaller dataset that included females who had phenotypic observations of annual breeding success as well as sequence data with which we could estimate F_GRM_ (see above). This dataset therefore included N_observations_ = 126 from N_females_ = 85.

## Results

### Temporal trend in precocial breeding

We did not find any evidence suggesting that annual breeding success changed through time for either adults or 1-year-olds (Table 1, Figure 1). Specifically, we did not find an effect of year on breeding success for either adults or 1-year-olds, irrespective of whether year was fit as a linear effect or as a smoothed term in the model (Table 1, Table S1 and Figure 2). Fitting year as a smoothed term in the model did appear to capture the initial increase in breeding success in 1-year-olds reported previously, however confidence in this was low and the posterior distribution of the predicted effect was very wide and not different from zero (Table 1, Figure 2).

**Figure 1.**
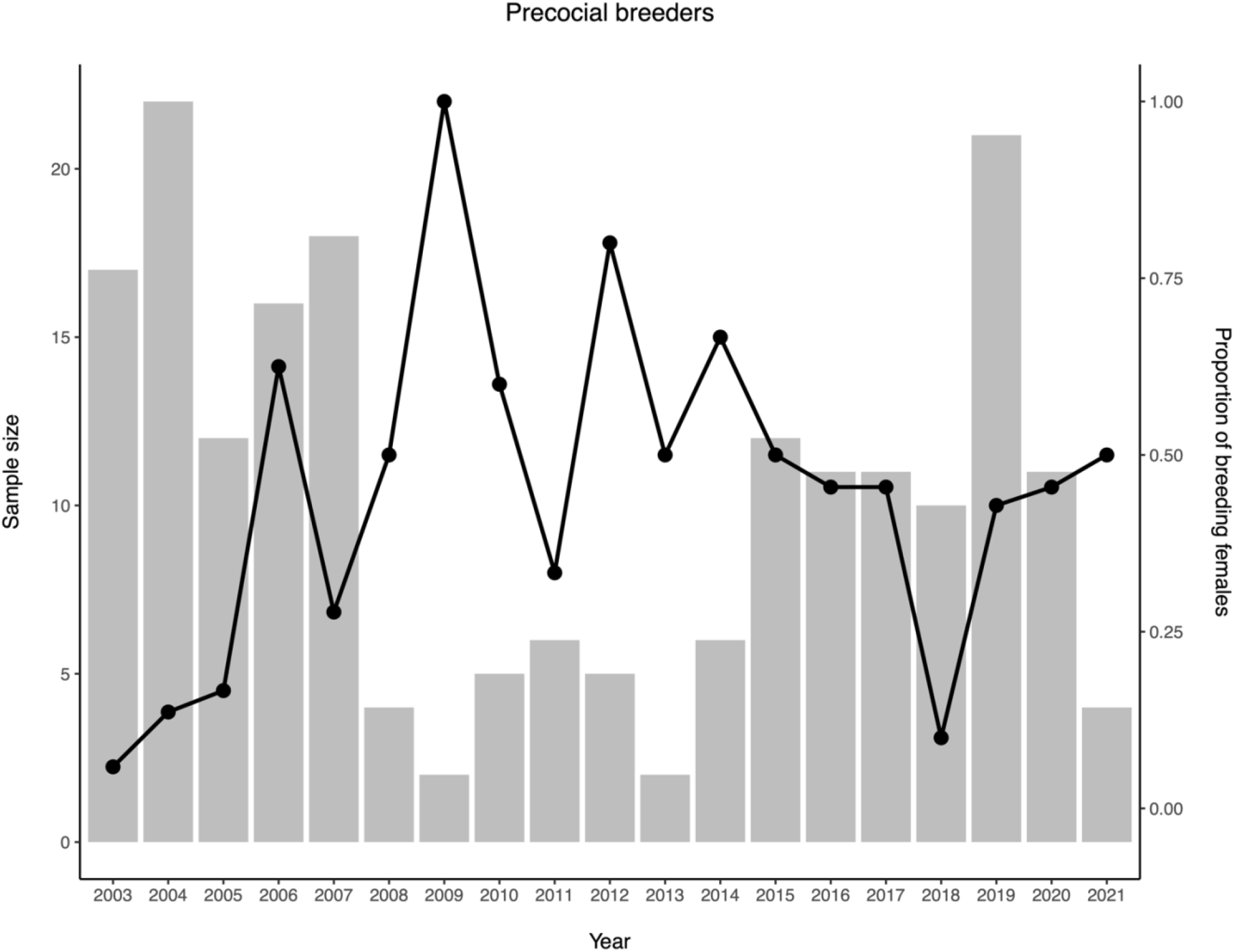
Plot showing the raw data for the number of one-year old females caught in each year (grey bars), and the proportion of those females that had bred (black points and line) for each year between 2003 and 2021.

**Figure 2.**
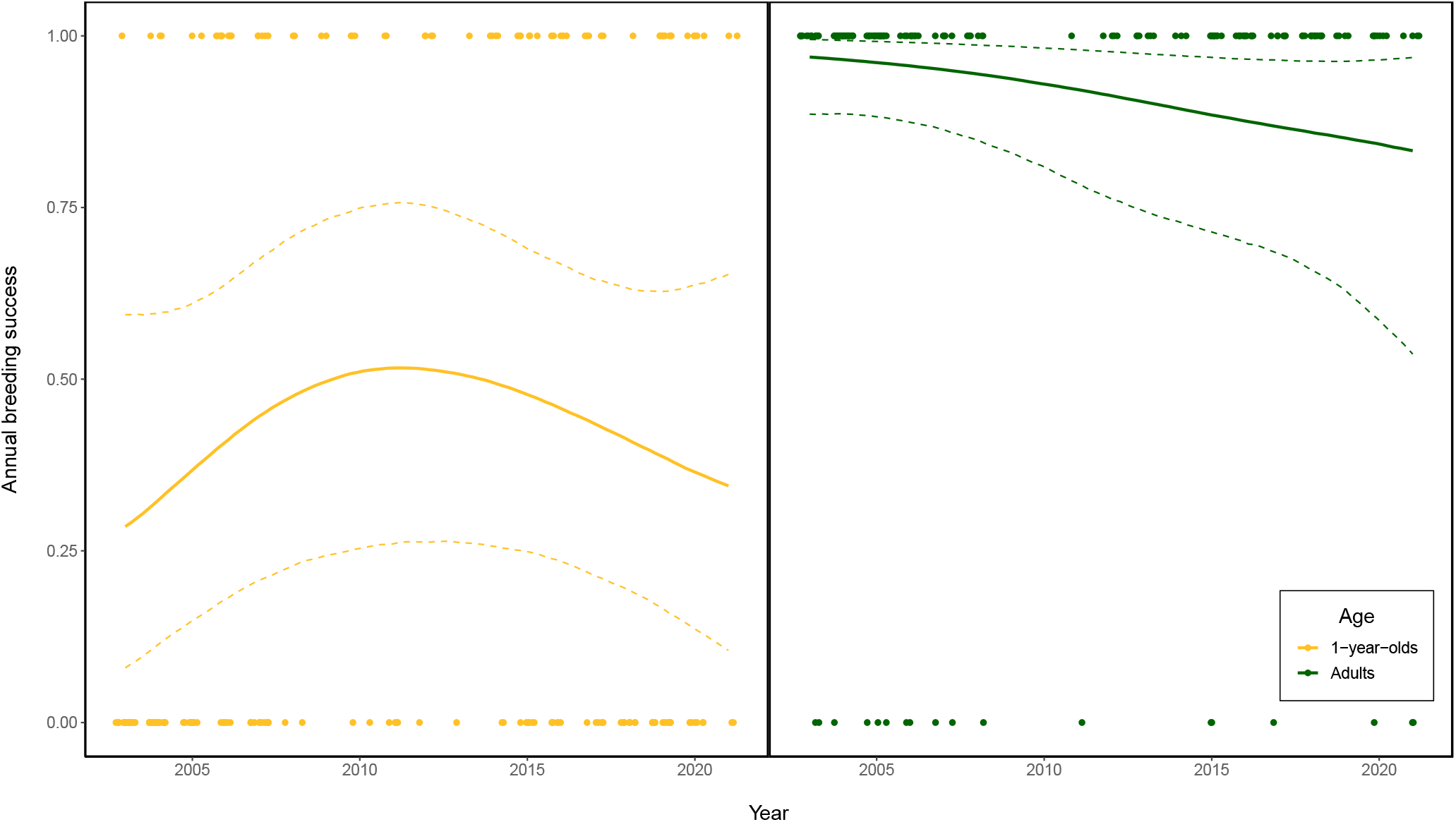
Plot showing the relationship between year and annual breeding success for 1-year- olds (gold) and adults (green). Points show observed data, and regression lines show the predicted relationship between year and annual breeding success derived from a mixed effects model which fits annual breeding success as a function of year, which was fit as a smoothed term with 5 knots (see methods for full model structure). Solid line shows predictions derived from the median of the posterior and the dotted lines show upper and lower 95% confidence intervals from posterior distribution.

**Table 1.** Table summarising results from a mixed effects model used to estimate relationship between size traits (weight and head width), time (year) and annual breeding success. Posterior medians of linear coefficient estimate for fixed effects and standard deviations for random effects presented with 95% credible intervals of posterior distribution in parentheses. Estimates where posterior does not overlap with zero in bold.

| Parameter |  |  |
| --- | --- | --- |
| Fixed Effects | Age<sub>ADULT</sub> | 0.03 (-9.67 - 9.79) |
|  | Year | 0.14 (-0.50 - 1.00) |
|  | Headwidth | 0.39 (-0.37 - 1.62) |
|  | Weight | **6.84 (2.02 - 13.53)** |
|  | DFTD | 2.39 (-3.33 - 8.55) |
|  | Age<sub>ADULT</sub>:Year | 0.08 (0.01 - 0.20) |
|  | Age<sub>ADULT</sub>:Headwidth | **-1.46 (-3.79 - -0.15)** |
|  | Age<sub>ADULT</sub>:Weight | 1.98 (-2.40 - 7.66) |
| Random effects | ID | 12.83 (3.18 - 28.16) |
|  | TrapID | 8.23 (1.68 - 18.86) |
|  | Year | 3.08 (0.18 - 8.71) |
|  | Month | 7.73 (1.08 - 20.98) |

### Relationship between breeding success and size

Head width did not have an effect on annual breeding success in female Tasmanian devils (Table 1, Figure 3). However, heavier females were more likely to breed than those with lower body weight (Table 1, Figure 3). We did not find evidence for an interaction between age and body weight (Table 3), suggesting that the pattern of heavier females being more likely to breed was consistent across 1-year olds and adults. We did find a significant interaction between head width and age (Table 1). However, this significant interaction seems unlikely to be biologically meaningful as the main effect for head width was not different from zero, and the cross-over effect that this interaction suggests does not appear until the larger values of head width measurements (Figure S1) where there are fewer data and therefore a great amount of uncertainty. There was no evidence for a relationship between having DFTD and likelihood of breeding in a given year (Table 1), suggesting that being infected with DFTD did not affect the probability of females breeding.

**Figure 3.**
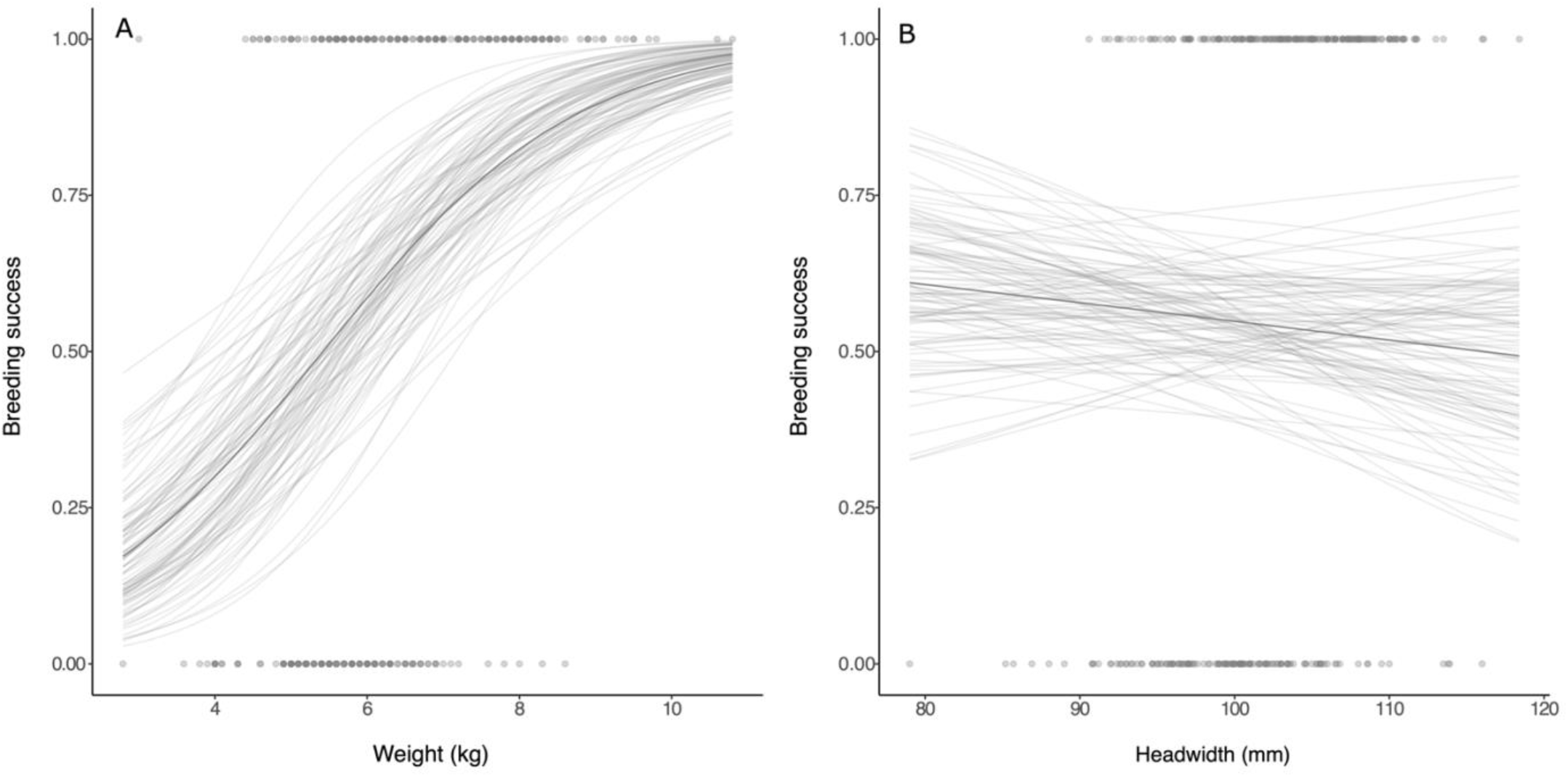
Plot showing the relationship between head width and annual breeding success (a), and weight with annual breeding success (b). Points show observed data, and regression lines show the predicted relationship between size traits and annual breeding success derived from a mixed effects model which fits annual breeding success as a function of both size traits (see methods for full model structure). Solid dark line shows predictions derived from the median of the posterior and the lighter lines show 100 randomly selected draws from the posterior.

### Inbreeding depression

We did not find evidence for a relationship between annual breeding success and the individual inbreeding coefficient, F_GRM,_ either for 1-year-olds of for adults (Table 2). This was true when we fit the full model, or fit a reduced model with just age and F_GRM_ fit as fixed effects, suggesting that removing the fixed effects with little statistical support in the previous model did not affect our estimates of the effect of inbreeding.

**Table 2.**
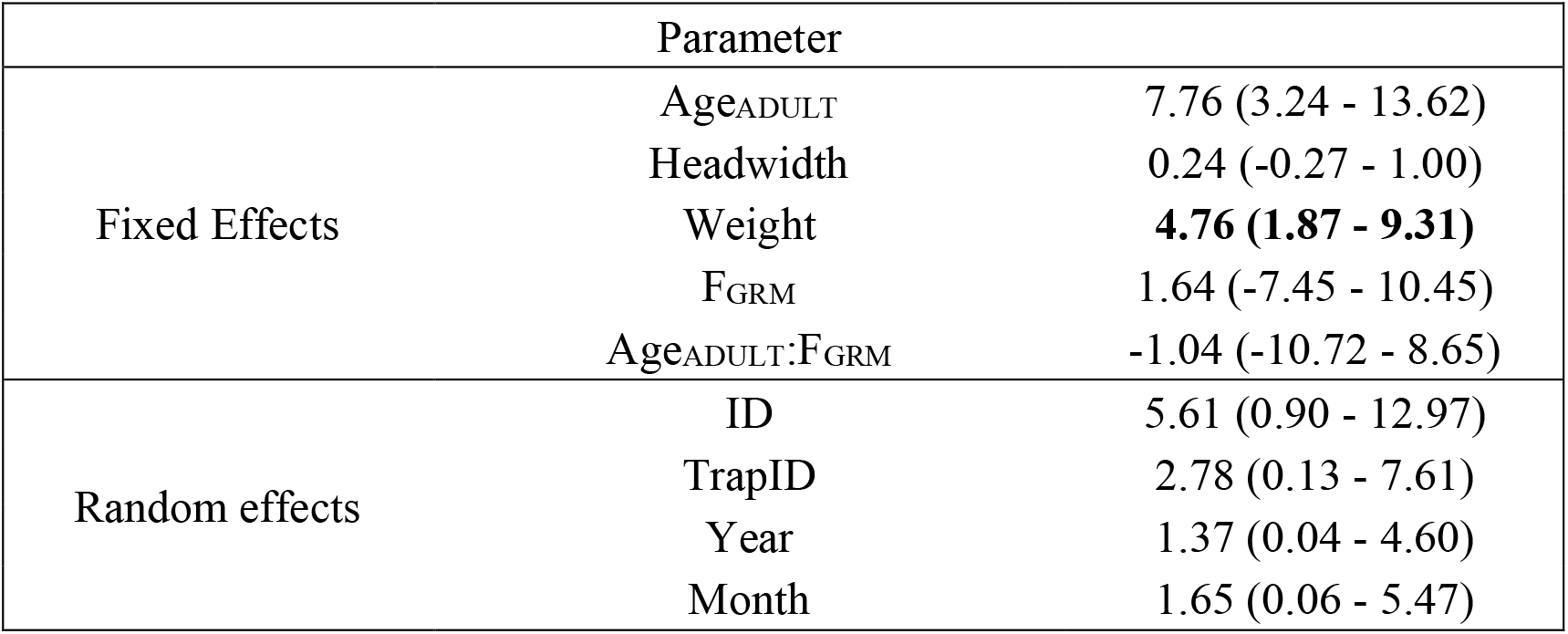
Table summarising results from a mixed effects model used to estimate inbreeding (F_GRM_) and annual breeding success. Posterior medians of linear coefficient estimates for fixed effects and standard deviations for random effects presented with 95% credible intervals of posterior distribution in parentheses. Estimates where posterior does not overlap with zero in bold.

| Parameter |  |  |
| --- | --- | --- |
| Fixed Effects | Age<sub>ADULT</sub> | 7.76 (3.24 - 13.62) |
|  | Headwidth | 0.24 (-0.27 - 1.00) |
|  | Weight | **4.76 (1.87 - 9.31)** |
|  | F<sub>GRM</sub> | 1.64 (-7.45 - 10.45) |
|  | Age<sub>ADULT</sub>:F<sub>GRM</sub> | -1.04 (-10.72 - 8.65) |
| Random effects | ID | 5.61 (0.90 - 12.97) |
|  | TrapID | 2.78 (0.13 - 7.61) |
|  | Year | 1.37 (0.04 - 4.60) |
|  | Month | 1.65 (0.06 - 5.47) |

## Discussion

We analysed data from a long term mark-recapture study of a wild, declining population of Tasmanian devils to test hypotheses related to observations of disease-induced female precocial breeding detected 20 years ago (15). We tested whether this life history shift, representing a classic trade-off between growth and reproduction, continued over time due to prior observations that age structure in populations with high DFTD prevalence collapsed (32) and that adult females (2 years old) became semelparous. Additionally, we tested for evidence of inbreeding depression, as population sizes collapsed by more than 80% in most infected devil populations. With 15 years of additional data, we found that despite an initial rise in rates of precocial breeding in the two-years following DFTD emergence, there was no further rise thereafter, and the probability of 1-year-olds breeding has plateaued at a rate of about 40% for the past 18 years. We also found that while heavier females were more likely to breed overall, there was no evidence for a differential effect of size on breeding success for 1-year-olds or individuals greater than two years old. Finally, we found that breeding success was not under inbreeding depression for either age category.

According to life-history theory, early reproduction should be selected for when population perturbances impact adult survival more than juvenile survival (40). DFTD almost exclusively impacts survival in adult Tasmanian devils, as the youngest a devil may contract the disease is around 14 months (15). The original rise in rates of precocial breeding found immediately after the emergence of DFTD (15) was probably in response to reduced intraspecific competition and enhanced resource availability following population decline (41). If precocial breeding provides a fitness benefit, we would expect that selection would favour increased juvenile growth rates or on smaller size at first breeding, either of which could result in an increased rate of precocial breeding over a period of 18 years. However, our analyses showed no significant interaction between female body mass and age, although heavier females are overall more likely to breed. These results suggest either that the initial response in precocial breeding reflects plasticity rather than selection, or that the population may have faced constraints that inhibited a further increase in proportion of precocial breeders. Such constraints could be caused, for instance, by limits imposed on individual growth rates in the population. That is, females that breed precocially normally breed in May which is late in the breeding season (28) and their female offspring cannot then grow enough to reach maturity by May upon weaning and den departure of the subsequent year, resulting in inhibition of precocial breeding. An additional constraint could be caused by having a mother with DFTD, which would compromise nutrition of her young as tumours grew (42). A promising avenue of future research would be to test this hypothesis once the necessary data are available.

Body weight is often positively correlated with reproductive success, due to enhanced ability to provision resources to offspring (43). Accordingly, body weight is often found to be under selection in populations of wild animals (44). Here, we found that heavier female Tasmanian devils were more likely to breed and this effect was not different between age classes. Thus, body weight may be under selection in this population, resulting in the expectation that and based on this result alone we might expect body weight should increase over time. Interestingly, this prediction is further supported by work showing that heavier individuals are also less likely to contract DFTD (45). However, in the Freycinet population (focal population here), mean body weight has not changed since 1999 (45). A lack of temporal change despite evidence for positive selection has been found in other systems (46,47) and may be due to a fluctuating environment that can alter the selective landscape experienced by the population which inhibits an overall positive evolutionary response (48–50). However, such a “paradox- of-stasis” remains a major unanswered question in evolutionary biology and empirical research is needed to test hypotheses that may explain how this phenomena emerges (51).

Inbreeding depression, defined as a negative correlation between heterozygosity and fitness (20), can be a serious risk to declining populations (18,21) and can affect life-stages differently (23–25). We found no evidence for inbreeding depression on breeding success in either adult or subadult females. These results are surprising in light of previous studies that show genetic diversity is quite low across the range of devils (52). A possible explanation is that deleterious alleles have been purged from the population during the observed population decline (53). Indeed, previous work has shown sufficient genetic diversity for devils to respond to selection (34,35,54–56), as well as a lack of evidence for inbreeding depression in susceptibility to DFTD (45). However, note that inbreeding depression is highly dependent on present and future environmental conditions, and may be present in other fitness-related traits not measured here (e.g., survival). Therefore, conclusions about the demographic consequences of inbreeding in Tasmanian devils should not be drawn from our results alone.

It is important to note that the lack of a relationship between inbreeding, size and breeding success reported here may not reflect patterns in Tasmanian devil populations outside the Freycinet peninsula. Moreover, these “null-results” should not be over-interpreted as the absence of an effect; they may have arisen instead as a result of lack of statistical power in our dataset. Nevertheless, our work shows that, despite an initial rise in proportion of females precocially breeding after the emergence of DFTD, the probability of 1-year-old females breeding has not continued to rise thereafter. Our work thus demonstrates the value of long-term, multigenerational population studies (57,58) and the importance of follow-up studies when trying to understand temporal changes in life-history traits. Finally, important questions remain about the underlying mechanisms associated with the initial rise of precocial breeding followed by a plateau observed here remain unknown. Determining the extent that those changes were associated with selection or plasticity, as well as tracking any temporal changes in growth rates, could ultimately provide key insights as to how life-history strategies may respond to population perturbances.

## Acknowledgements

We would like to first acknowledge the Toorerno-maire-mener clan, the Traditional Custodians and First Peoples of the Freycinet Peninsula where this project was delivered, and we pay respect to their Elders past, present and emerging. We would also like to thank the many researchers involved in field sampling during the course of the study. We also thank Soraia Barbosa for facilitating the sharing of the genetic data.

## References

1. Stearns SC. Trade-Offs in Life-History Evolution. Funct Ecol. 1989;3(3):259–68.

2. Roff. Trade-offs between growth and reproduction: an analysis of the quantitative genetic evidence. J Evol Biol. 2000 May 1;13(3):434–45.

3. Roff DA, Fairbairn DJ. The evolution of trade-offs: where are we? J Evol Biol. 2007 Mar 1;20(2):433–47.

4. Butler SJ, Benton TG, Nicoll MAC, Jones CG, Norris K. Indirect Population Dynamic Benefits of Altered Life-History Trade-Offs in Response to Egg Harvesting. Am Nat. 2009 Jul 1;174(1):111–21.

5. De Roos AM, Persson L, McCauley E. The influence of size-dependent life-history traits on the structure and dynamics of populations and communities. Ecol Lett. 2003 May 1;6(5):473–87.

6. Fowler CW. Density Dependence as Related to Life History Strategy. Ecology. 1981 Jun 1;62(3):602–10.

7. Baruah G, Clements CF, Guillaume F, Ozgul A. When Do Shifts in Trait Dynamics Precede Population Declines? Am Nat. 2019 May 1;193(5):633–44.

8. Forcada J, Trathan PN, Murphy EJ. Life history buffering in Antarctic mammals and birds against changing patterns of climate and environmental variation. Glob Change Biol. 2008 Nov 1;14(11):2473–88.

9. Snell-Rood E, Cothran R, Espeset A, Jeyasingh P, Hobbie S, Morehouse NI. Life-history evolution in the anthropocene: effects of increasing nutrients on traits and trade-offs. Evol Appl. 2015 Aug 1;8(7):635–49.

10. Bronikowski AM, Clark ME, Rodd FH, Reznick DN. POPULATION-DYNAMIC CONSEQUENCES OF PREDATOR-INDUCED LIFE HISTORY VARIATION IN THE GUPPY (POECILIA RETICULATA). Ecology. 2002 Aug 1;83(8):2194–204.

11. Claessen D, Van Oss C, de Roos AM, Persson L. THE IMPACT OF SIZE-DEPENDENT PREDATION ON POPULATION DYNAMICS AND INDIVIDUAL LIFE HISTORY. Ecology. 2002 Jun 1;83(6):1660–75.

12. Day T, Abrams PA, Chase JM. THE ROLE OF SIZE-SPECIFIC PREDATION IN THE EVOLUTION AND DIVERSIFICATION OF PREY LIFE HISTORIES. Evolution. 2002 May 1;56(5):877–87.

13. Jørgensen C, Ernande B, Fiksen Ø. Size-selective fishing gear and life history evolution in the Northeast Arctic cod. Evol Appl. 2009 Aug 1;2(3):356–70.

14. Olsen EM, Heino M, Lilly GR, Morgan MJ, Brattey J, Ernande B, et al. Maturation trends indicative of rapid evolution preceded the collapse of northern cod. Nature. 2004 Apr 1;428(6986):932–5.

15. Jones ME, Cockburn A, Hamede R, Hawkins C, Hesterman H, Lachish S, et al. Lifehistory change in disease-ravaged Tasmanian devil populations. Proc Natl Acad Sci. 2008 Jul 22;105(29):10023–7.

16. Valenzuela-Sánchez A, Wilber MQ, Canessa S, Bacigalupe LD, Muths E, Schmidt BR, et al. Why disease ecology needs life-history theory: a host perspective. Ecol Lett. 2021 Apr 1;24(4):876–90.

17. Capdevila P, Stott I, Cant J, Beger M, Rowlands G, Grace M, et al. Life history mediates the trade-offs among different components of demographic resilience. Ecol Lett. 2022 Jun 1;25(6):1566–79.

18. O’Grady JJ, Brook BW, Reed DH, Ballou JD, Tonkyn DW, Frankham R. Realistic levels of inbreeding depression strongly affect extinction risk in wild populations. Biol Conserv. 2006 Nov 1;133(1):42–51.

19. Charlesworth B, Hughes KA. Age-specific inbreeding depression and components of genetic variance in relation to the evolution of senescence. Proc Natl Acad Sci. 1996 Jun 11;93(12):6140–5.

20. Charlesworth D, Willis JH. The genetics of inbreeding depression. Nat Rev Genet. 2009 Nov 1;10(11):783–96.

21. Hedrick PW, Kalinowski ST. Inbreeding Depression in Conservation Biology. Annu Rev Ecol Syst. 2000 Nov 1;31(1):139–62.

22. Chen N, Cosgrove EJ, Bowman R, Fitzpatrick JW, Clark AG. Genomic Consequences of Population Decline in the Endangered Florida Scrub-Jay. Curr Biol. 2016 Nov 7;26(21):2974–9.

23. Huisman J, Kruuk LEB, Ellis PA, Clutton-Brock T, Pemberton JM. Inbreeding depression across the lifespan in a wild mammal population. Proc Natl Acad Sci. 2016 Mar 29;113(13):3585–90.

24. Stoffel MA, Johnston SE, Pilkington JG, Pemberton JM. Genetic architecture and lifetime dynamics of inbreeding depression in a wild mammal. Nat Commun. 2021 May 20;12(1):2972.

25. Keller LF, Reid JM, Arcese P. Testing evolutionary models of senescence in a natural population: age and inbreeding effects on fitness components in song sparrows. Proc R Soc B Biol Sci. 2008 Jan 23;275(1635):597–604.

26. Cunningham CX, Comte S, McCallum H, Hamilton DG, Hamede R, Storfer A, et al. Quantifying 25 years of disease-caused declines in Tasmanian devil populations: host density drives spatial pathogen spread. Ecol Lett. 2021 May 1;24(5):958–69.

27. McCallum H, Tompkins DM, Jones M, Lachish S, Marvanek S, Lazenby B, et al. Distribution and Impacts of Tasmanian Devil Facial Tumor Disease. EcoHealth. 2007 Sep 1;4(3):318–25.

28. Lachish S, McCallum H, Jones M. Demography, Disease and the Devil: Life-History Changes in a Disease-Affected Population of Tasmanian Devils (Sarcophilus harrisii). J Anim Ecol. 2009;78(2):427–36.

29. Chang C chen, Moiron M, Sánchez-Tójar A, Niemelä PT, Laskowski KL. What is the meta-analytic evidence for life-history trade-offs at the genetic level? Ecol Lett. 2024 Jan 1;27(1):e14354.

30. Lachish S, Jones M, McCallum H. The Impact of Disease on the Survival and Population Growth Rate of the Tasmanian Devil. J Anim Ecol. 2007;76(5):926–36.

31. Jones ME. Over-eruption in marsupial carnivore teeth: compensation for a constraint. Proc R Soc B Biol Sci. 2023 Dec 13;290(2013):20230644.

32. Hamede RK, Pearse AM, Swift K, Barmuta LA, Murchison EP, Jones ME. Transmissible cancer in Tasmanian devils: localized lineage replacement and host population response. Proc R Soc B Biol Sci. 2015 Sep 7;282(1814):20151468.

33. Hesterman H, Jones SM, Schwarzenberger F. Pouch appearance is a reliable indicator of the reproductive status in the Tasmanian devil and the spotted-tailed quoll. J Zool. 2008 Jun 1;275(2):130–8.

34. Epstein B, Jones M, Hamede R, Hendricks S, McCallum H, Murchison EP, et al. Rapid evolutionary response to a transmissible cancer in Tasmanian devils. Nat Commun. 2016 Aug 30;7(1):12684.

35. Margres MJ, Jones ME, Epstein B, Kerlin DH, Comte S, Fox S, et al. Large-effect loci affect survival in Tasmanian devils (Sarcophilus harrisii) infected with a transmissible cancer. Mol Ecol. 2018 Nov 1;27(21):4189–99.

36. Ali OA, O’Rourke SM, Amish SJ, Meek MH, Luikart G, Jeffres C, et al. RAD Capture (Rapture): Flexible and Efficient Sequence-Based Genotyping. Genetics. 2016 Feb 1;202(2):389–400.

37. Murchison EP, Schulz-Trieglaff OB, Ning Z, Alexandrov LB, Bauer MJ, Fu B, et al. Genome Sequencing and Analysis of the Tasmanian Devil and Its Transmissible Cancer. Cell. 2012 Feb 17;148(4):780–91.

38. Yang J, Lee SH, Goddard ME, Visscher PM. GCTA: a tool for genome-wide complex trait analysis. Am J Hum Genet. 2011 Jan 7;88(1):76–82.

39. Bürkner PC. brms: An R package for Bayesian multilevel models using Stan. J Stat Softw. 2017;80(1):1–28.

40. Charnov EL, Schaffer WM. Life-History Consequences of Natural Selection: Cole’s Result Revisited. Am Nat. 1973 Nov 1;107(958):791–3.

41. Comte S, Carver S, Hamede R, Jones M. Changes in spatial organization following an acute epizootic: Tasmanian devils and their transmissible cancer. Glob Ecol Conserv. 2020 Jun 1;22:e00993.

42. Ruiz-Aravena M, Jones ME, Carver S, Estay S, Espejo C, Storfer A, et al. Sex bias in ability to cope with cancer: Tasmanian devils and facial tumour disease. Proc R Soc B Biol Sci. 2018 Nov 21;285(1891):20182239.

43. Blanckenhorn WU. The Evolution of Body Size: What Keeps Organisms Small? Q Rev Biol. 2000 Dec 1;75(4):385–407.

44. Kingsolver JG, Pfennig DW. INDIVIDUAL-LEVEL SELECTION AS A CAUSE OF COPE’S RULE OF PHYLETIC SIZE INCREASE. Evolution. 2004 Jul 1;58(7):1608–12.

45. Strickland K, Jones ME, Storfer A, Hamede RK, Hohenlohe PA, Margres MJ, et al. Adaptive potential in the face of a transmissible cancer in Tasmanian devils. Mol Ecol. 2024 Sep 28;n/a(n/a):e17531.

46. Gotanda KM, Correa C, Turcotte MM, Rolshausen G, Hendry AP. Linking macrotrends and microrates: Re-evaluating microevolutionary support for Cope’s rule. Evolution. 2015 May 1;69(5):1345–54.

47. O’Sullivan RJ, Aykanat T, Johnston SE, Kane A, Poole R, Rogan G, et al. Evolutionary stasis of a heritable morphological trait in a wild fish population despite apparent directional selection. Ecol Evol. 2019 Jun 1;9(12):7096–111.

48. Bell G. Fluctuating selection: the perpetual renewal of adaptation in variable environments. Philos Trans R Soc B Biol Sci. 2010 Jan 12;365(1537):87–97.

49. de Villemereuil P, Charmantier A, Arlt D, Bize P, Brekke P, Brouwer L, et al. Fluctuating optimum and temporally variable selection on breeding date in birds and mammals. Proc Natl Acad Sci. 2020 Dec 15;117(50):31969–78.

50. Wright J, Bolstad GH, Araya-Ajoy YG, Dingemanse NJ. Life-history evolution under fluctuating density-dependent selection and the adaptive alignment of pace-of-life syndromes. Biol Rev. 2019 Feb 1;94(1):230–47.

51. Merilä J, Sheldon BC, Kruuk LEB. Explaining stasis: microevolutionary studies in natural populations. Genetica. 2001 Nov 1;112(1):199–222.

52. Jones ME, Paetkau D, Geffen E, Moritz C. Genetic diversity and population structure of Tasmanian devils, the largest marsupial carnivore. Mol Ecol. 2004 Aug 1;13(8):2197– 209.

53. Hedrick PW, Garcia-Dorado A. Understanding Inbreeding Depression, Purging, and Genetic Rescue. Trends Ecol Evol. 2016 Dec 1;31(12):940–52.

54. Fraik AK, Margres MJ, Epstein B, Barbosa S, Jones M, Hendricks S, et al. Disease swamps molecular signatures of genetic-environmental associations to abiotic factors in Tasmanian devil (Sarcophilus harrisii) populations. Evolution. 2020 Jul 1;74(7):1392– 408.

55. Stahlke AR, Epstein B, Barbosa S, Margres MJ, Patton AH, Hendricks SA, et al. Contemporary and historical selection in Tasmanian devils (Sarcophilus harrisii) support novel, polygenic response to transmissible cancer. Proc R Soc B Biol Sci. 2021 May 26;288(1951):20210577.

56. Gallinson DG, Kozakiewicz CP, Rautsaw RM, Beer MA, Ruiz-Aravena M, Comte S, et al. Intergenomic signatures of coevolution between Tasmanian devils and an infectious cancer. Proc Natl Acad Sci. 2024 Mar 19;121(12):e2307780121.

57. Clutton-Brock TH, Sheldon BC. Individuals and populations: the role of long-term, individual-based studies of animals in ecology and evolutionary biology. Trends Ecol Evol. 2010;25(10):562–73.

58. Sheldon BC, Kruuk LEB, Alberts SC. The expanding value of long-term studies of individuals in the wild. Nat Ecol Evol. 2022 Dec 1;6(12):1799–801.

