## Supplemental Table 1 for "Effects of a transmissible cancer on life-history traits in Tasmanian devils"

### Supplementary material

**Table S1.** Table summarising results from a mixed effects model used to estimate relationship between size traits (weight and headwidth), time (year) and annual breeding success. Year was fit as a smoothed, non-linear term with five knots. Posterior medians of linear coefficient estimates for fixed effects and standard deviations for random effects presented with 95% credible intervals of posterior distribution in parentheses. Estimates where posterior does not overlap with zero in bold.

| | Parameter | |
|---|---|---|
| Fixed Effects | AgeADULT | 3.81 (-5.95 - 13.30) |
| | Headwidth | -0.07 (-0.46 - 0.18) |
| | Weight | **2.72 (0.66 - 8.28)** |
| | DFTD | 1.52 (-0.76 - 5.72) |
| | AgeADULT:Headwidth | -0.02 (-0.18 - 0.21) |
| | AgeADULT:Weight | 1.05 (-0.97 - 4.06) |
| | sAgeADULT:Year | -3.88 (-11.60 - 4.34) |
| | sAge1yearold:Year | 2.41 (-4.85 - 10.09) |
| Random Effects | ID | 4.08 (0.15 - 13.76) |
| | TrapID | 2.37 (0.09 - 8.43) |
| | Year | 1.48 (0.11 - 4.97) |
| | Month | 2.76 (0.16 - 10.09) |
